# Trunk muscle activity across various trunk–limb coordination patterns in centipede locomotion

**DOI:** 10.64898/2026.11.05.764723

**Authors:** Daiki Wakita, Kotaro Yasui

## Abstract

Animals coordinate movements of their limbs and flexible bodies, yet how trunk muscle activity changes across locomotor patterns remains unclear. We simultaneously recorded electromyograms from trunk muscles (deep dorsal oblique muscles) and trunk–limb kinematics in two centipede species exhibiting retrograde and direct locomotor waves during swimming and walking. During swimming, trunk muscle activity followed a similar phase relationship with trunk bending in both species despite opposite wave directions and different involvement of limbs, suggesting common active generation of undulation. During fast walking, trunk muscle activity was strongest around the ipsilateral swing phase in the undulatory retrograde-wave gait of *Scolopendra mutilans*, but weaker and associated with the ipsilateral stance phase despite the nearly straight trunk in the direct-wave gait of *Scolopocryptops rubiginosus*. Together, our findings suggest that the same trunk muscles serve two contrasting mechanical roles depending on locomotor mode: either generating trunk undulation or actively straightening the trunk against limbinduced deformation.

## INTRODUCTION

Animal locomotion emerges from interactions among body morphology, neuromuscular control, and the physical environment, giving rise to a wide variety of movement patterns even within closely related taxa. Understanding how animals generate and switch between different locomotor patterns remains a central problem in biomechanics and neuroethology. Myriapods are particularly suited to this question because their long, flexible trunks and numerous limbs enable various locomotor patterns.

Centipedes exhibit locomotor patterns involving rhythmic trunk bending, coordinated movements of numerous limbs, or both (Manton, 1952; Lewis, 1981). Their locomotion has been classified into two major gait patterns, retrograde-wave and direct-wave gaits, which differ in the direction of metachronal waves of trunk segments and/or limbs (Fig. 1). In retrograde-wave locomotion, the metachronal waves propagate from anterior to posterior, and limbs on the same side tend to share similar footholds (Fig. 1A). In contrast, direct-wave locomotion is characterized by non-overlapping footholds associated with metachronal waves propagating from posterior to anterior (Fig. 1B). The direction of metachronal waves and the extent of trunk bending vary depending on species, locomotion speed, and substrates (Manton, 1952; Lewis, 1981; Kuroda et al., 2022). For example, *Scolopendra mutilans* exhibits a trunk-bending wave that is coordinated with the retrograde limb wave as its walking speed increases (Fig. 1C). However, during locomotion on the water surface, it folds limbs against the body and propels itself exclusively through the retrograde undulation of the trunk (Fig. 1D; Yasui et al., 2019). Another species, the scolopendromorph *Scolopocryptops rubiginosus*, displays a different locomotor strategy; it exhibits retrograde limb waves at low speeds and shifts to direct limb waves at high speeds, while maintaining a nearly straight trunk in both states (Fig. 1A, B; Kuroda et al., 2022). When moving on the water surface, this species employs direct trunk-bending waves that are coordinated with the direct limb waves (Fig. 1E; Kuroda et al., 2022). To date, these gait differences have been described at the kinematic level, whereas little is known about the underlying motor activity that produces them.

**Figure 1.**
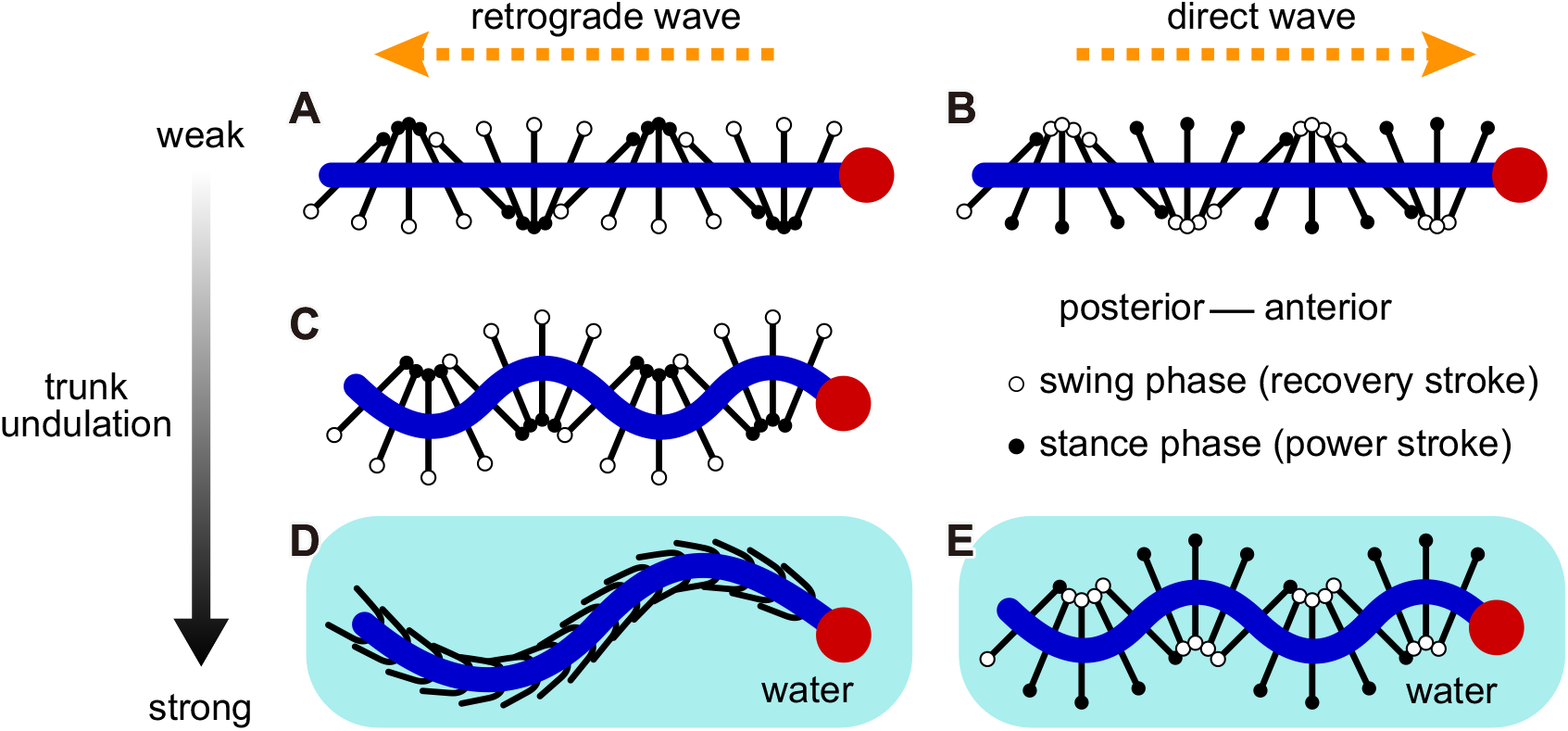
Locomotor modes of centipedes. (A) Retrograde-wave gait with small trunk undulation. (B) Direct-wave gait with small trunk undulation. (C) Retrograde-wave gait with large trunk undulation. (D) Retrograde-wave gait with large trunk undulation and folded limbs on the water surface. (E) Direct-wave gait with large trunk undulation on the water surface. In (A–C,E), the left and right limbs are assumed to move in antiphase.

Electromyography provides a means to reveal the motor control that cannot be inferred from kinematics alone. Anderson et al. (1995) investigated muscle activity during terrestrial locomotion in *Scolopendra heros*; however, their analysis was limited to a single species exhibiting only a retrograde-wave pattern and did not consider locomotion on the water surface. Consequently, how muscle activation varies across different gait patterns, including direct-wave and water-surface locomotion, remains unknown. In this study, we address this gap by simultaneously recording trunk and limb kinematics together with the electromyograms (EMGs) of trunk muscles, and by comparing retrograde-versus direct-wave locomotion as well as terrestrial versus water-surface locomotion in two centipede species, *S. mutilans* and *S. rubiginosus* (Fig. 2A,B).

**Figure 2.**
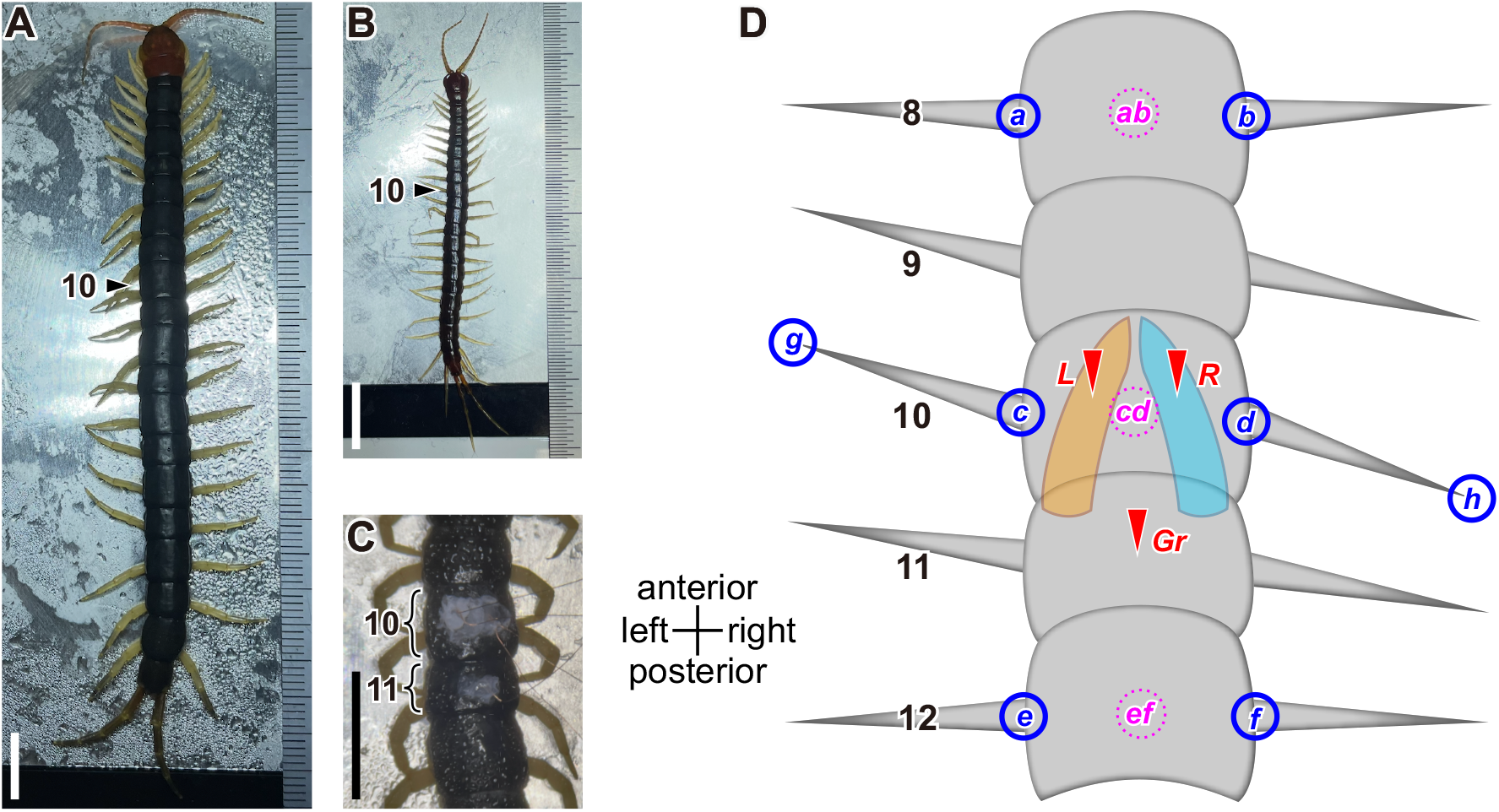
Animals and schematic of kinematic and electromyogram (EMG) measurement. (A) *Scolopendra mutilans*. (B) *Scolopocryptops rubiginosus*. (C) Magnified view of the region around trunk segment 10 after electrode-wire insertion in *S. mutilans*. The whitish material is cyanoacrylate adhesive. In (A–C), the animals were cooled on a plate for electrode insertion. (D) Tracking points and electrode-insertion sites. Solid circles represent the tracking points at the limb bases (*a*–*f*) and limb tips (*g, h*). Dotted circles (*ab, cd, ef*) represent the midpoints of left and right limb bases in the same segments. Arrowheads indicate the sites of electrode insertion: *L*, left deep dorsal oblique muscle (DDOM); *R*, right DDOM; *Gr*, ground electrode site. In each panel, numbers on the left indicate the identification numbers of limb-bearing segments. Scale bars in (A–C): 1 cm.

## METHODS

### Animals

Adult *Scolopendra mutilans* (Chinese red-headed centipedes, Fig. 2A) were collected in Wakayama Prefecture and adult *Scolopocryptops rubiginosus* (Fig. 2B) in Miyagi Prefecture, Japan. Body lengths excluding antennae and ultimate legs were approximately 10–11 cm in *S. mutilans* and 4.5–5 cm in *S. rubiginosus*. Each individual was kept separately in a plastic container with a moistened paper towel at 23^*°*^C and fed insect jelly (Pro Jelly, KB Farm, Saitama, Japan) once per week.

### Electromyography

To examine trunk muscle activity across locomotor modes, we targeted the deep dorsal oblique muscles (DDOMs), whose anatomical arrangement is suited to lateral trunk flexion (Manton, 1965; Anderson et al., 1995). We selected the 10th trunk segment because it is a relatively long segment near the middle of the body (Fig. 2A–C), to minimize potential influences of the anterior and posterior body ends.

Bipolar wire electrodes were made from 0.05-mm-diameter enamel-coated copper wire (Kyowa Harmonet, Tokyo, Japan). Approximately 0.2 mm of insulation was removed from the wire tip, which was then bent at a right angle. Animals were anesthetized on a cooled plate at 1^*°*^C (SCP-125, AS ONE, Osaka, Japan) (Fig. 2A,B), and small openings were made on the left and right parts of the tergite of the 10th trunk segment using an ultrasonic cutter (ZO-41 II, Honda Electronics, Aichi, Japan). Through each opening, a recording electrode and a reference electrode were inserted presumably into the DDOM (Fig. 2D, arrowheads *L* and *R*). A ground electrode was inserted at the middle of the 11th-segment tergite (Fig. 2D, arrowhead *Gr*). Electrodes were secured with cyanoacrylate adhesive (Fig. 2C) and bundled using a small piece of aluminum foil.

Individuals were placed in an acrylic chamber inside a Faraday cage. Electrode tension was adjusted so that the animals exhibited locomotor patterns without forward displacement, allowing stable EMG recordings in a limited space. For swimming experiments, the chamber was filled with tap water, enabling the animals to float and swim on the water surface. Locomotor behavior was recorded from above (dorsal view), using a high-speed camera (HAS-U2, DITECT, Tokyo, Japan) at 200 fps. EMG signals were sampled at 1 kHz, amplified using an amplifier (0.1 kHz high-pass filter; DAM80, World Precision Instruments, Sarasota, FL, USA), and visualized with data-acquisition software (HAS-X Viewer; DITECT, Tokyo, Japan).

### Tracking

To relate trunk and limb kinematics to the EMG recordings, we focused tracking on the region around the 10th trunk segment. A 2-s period with stable kinematic and EMG patterns was selected for each trial. Trunk and limb kinematics were tracked using the software Tracker (Brown et al., 2025) at 100 fps. Tracked points were:

- the positions of the left and right limb bases of the 8th, 10th, and 12th trunk segments, each defined as the boundary between the tergite and the limb (circles *a* to *f* in Fig. 2D), and
- the positions of the distal tips of the left and right limbs of the 10th segment (circles *g* and *h*).

#### Trunk angle

Trunk bending was quantified as a body angle centered at the 10th segment. The midpoint between the left and right limb bases was calculated for each of the three segments (dotted circles *ab, cd*, and *ef* in Fig. 2D). Two vectors were then defined: one from the midpoint of segment 10 to that of 8 (vector *ab*–*cd*), and the other from the midpoint of segment 12 to that of 10 (vector *ef* –*cd*). The signed angle between these vectors was calculated, yielding a measure of local trunk curvature at segment 10. Positive values indicate that the segmented midline across segments 8, 10, and 12 (segment *ab*–*cd*–*ef*) is convex to the left, and negative values indicate convexity to the right.

#### Limb angle

Limb movements were quantified as signed angles of the left and right limbs of segment 10 relative to the trunk. Specifically, the limb angle was defined as the angle of a vector from the limb base to its distal tip (vectors *c*–*g* and *d*–*h* in Fig. 2D), measured relative to the left-right axis defined by the line connecting the bases of the right and left limbs of segment 10 (vector *d*–*c*). An angle of 0 indicates that the limb was oriented laterally, positive values indicate anterior orientation, and negative values indicate posterior orientation.

### Kinematic phase definition

To examine coordination among trunk and limb movements and the timing of EMG activity relative to these movements, we extracted kinematic phases from trunk and limb angle traces (artificial data shown in Fig. S1). Before phase extraction, trunk and limb angle traces were mean-centered within each trial and smoothed using a 51-ms running mean filter.

#### Trunk-based phase

When the trunk angle was used to define the locomotor phase, EMG activity was mapped to the phase of trunk bending relative to the EMG recording side (Fig. 3A). We defined the phases as follows:

- phases 0 and *π* as zero crossings (approximately straight posture),
- *π*/2 as maximal trunk bending toward the lateral side where the EMG was recorded (convex bending), and
- 3*π*/2 as maximal bending away from that side (concave bending).

**Figure 3.**
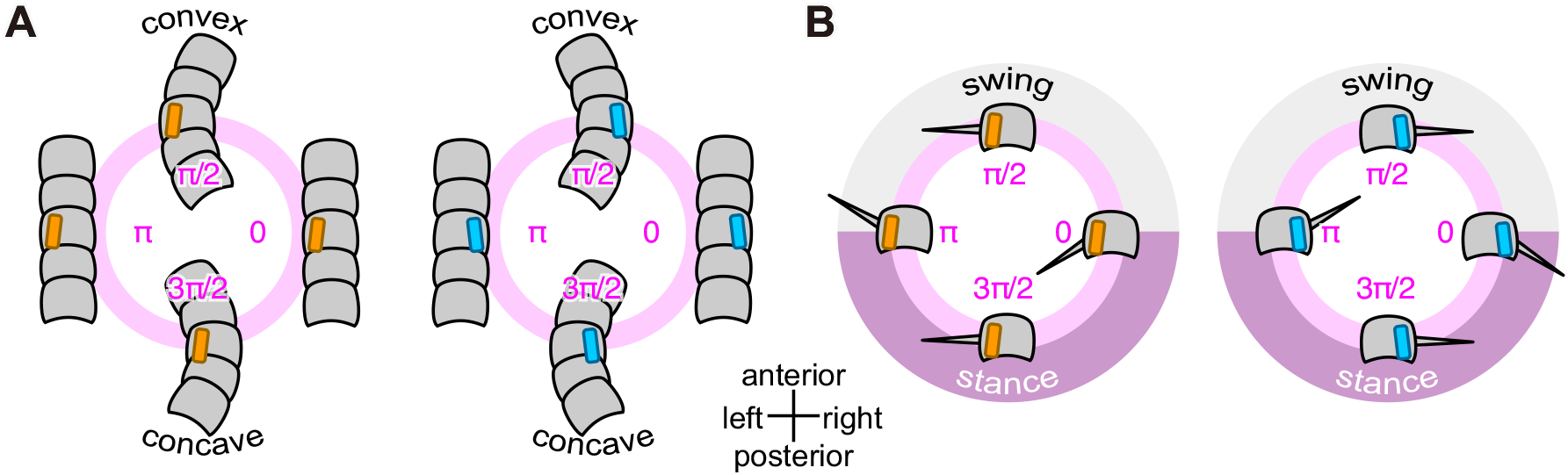
Kinematic phase definitions. In each panel, definitions for left and right sides are shown. For calculating trunk–limb phase differences, the trunk and limb phases were defined with reference to the same lateral side (Fig. 7A,B). For mapping electromyogram (EMG) intensity onto kinematic phases, the kinematic phase on the EMG recording side was used (Fig. 8). (A) Trunk-based phase: *π*/2 and 3*π*/2 correspond to maximal convex and concave bending relative to the reference side, respectively. (B) Limb-based phase: 0 and *π* correspond to the maximally posterior and anterior limb positions, respectively, for both left and right limbs.

In the trunk-angle trace, phase 0 corresponds to the upward zero crossing for left-side recordings, but to the downward zero crossing for right-side recordings. Phases other than 0, *π*/2, *π*, and 3*π*/2 were assigned to each time point by linear interpolation, yielding a continuous phase variable ranging from 0 to 2*π* for each cycle.

#### Limb-based phase

When the limb angles were used to define the locomotor phases, EMG activity was mapped to the phase of the limb on the EMG recording side (Fig. 3B). We defined the phases as follows:

- the negative peak as phase 0 (maximally posterior),
- the upward zero crossing as *π*/2 (mid forward),
- the positive peak as *π* (maximally anterior), and
- the downward zero crossing as 3*π*/2 (mid backward).

Phases 0 and *π* approximately correspond to the stance-to-swing and swing-to-stance transitions, respectively, although stance and swing were inferred from dorsal-view kinematics. As in the trunk-based phase analysis, intermediate phases were assigned by linear interpolation.

### Kinematic analysis

To characterize basic locomotor and coordination patterns before examining their relationships with EMG activity, we quantified trunk angular range and limb duty ratio and examined phase relationships among kinematic variables.

#### Trunk angular range

For each 2-s trial, the angular range of trunk undulation was calculated as the difference between the maximum and minimum values of the smoothed trunk angle.

#### Limb duty ratio

For each limb cycle, the duty ratio was calculated as the duration from phase *π* to the subsequent phase 2*π* (equivalent to 0), divided by the duration between consecutive occurrences of phase *π*. Thus, the duty ratio represents the fraction of one locomotor cycle from the maximally anterior limb position to the subsequent maximally posterior position, corresponding approximately to the stance phase as inferred from dorsal-view kinematics.

#### Phase difference

Because phase differences between body parts can vary within a locomotor cycle, we quantified them at four representative phases (0, *π/*2, *π*, and 3*π/*2) of a reference body part. The phase difference was calculated by subtracting the phase of the reference body part from that of the other body part and wrapping the resulting angle to the range 0–2*π*. Phase differences at each representative phase were summarized using the circular mean and phase consistency (mean resultant length).

### EMG analysis

EMG recordings were visually inspected for recording quality before phase analysis. Recordings were excluded when they showed pronounced low-frequency baseline fluctuations, indicative of movement artifacts, or when no discernible EMG activity above background noise was detected despite ongoing locomotion.

EMG intensity traces from the left and right DDOMs were analyzed at the original sampling rate. Each raw EMG signal was median-centered within each trial, and its absolute value was then taken to obtain a positive-valued EMG intensity trace (artificial data shown in Fig. S1). To reduce variability in recording amplitude among individuals and recording sites, EMG intensity was normalized separately for each individual and recording side. Mean EMG intensity was calculated for each locomotor mode by pooling all frames across trials, and all EMG intensity values were then divided by the largest of these mode-specific means. This procedure preserved relative differences in EMG intensity among locomotor modes.

To examine the relationships between the trunk muscle activity and the trunk/limb kinematics, EMG intensity was mapped onto the kinematic phase of the trunk and limbs (artificial data shown in Fig. S1). Each locomotor cycle was determined as the period between consecutive upward zero crossings (negative to positive transitions). We visualized relationships between kinematic phases and EMG intensity using circular plots. Phase values were binned in intervals of *π/*36, and EMG intensity was averaged within each bin.

To quantify how muscle activity depends on kinematic phase, we fitted a cosinor model to the relationship between kinematic phase *ϕ* and normalized EMG intensity *I*, namely, *I* = *α* + *β* cos*ϕ* + *γ* sin*ϕ*. Parameters were estimated by ordinary least-squares regression. The preferred phase of EMG activity *µ* and modulation amplitude *A* were calculated from the fitted coefficients as *µ* = atan2(*γ, β*) and 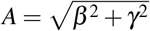.

## RESULTS

### Locomotor mode

Kinematic data and EMG recordings were obtained from multiple individuals of *Scolopendra mutilans* and *Scolopocryptops rubiginosus* across three locomotor conditions, i.e., swimming, fast walking, and slow walking (Table 1, Fig. 4, Video S1). In *S. mutilans*, all three locomotor modes showed retrograde waves. During swimming, the trunk exhibited strong undulation, with an angular range of 60 ± 8 deg (mean ± SD, Figs 5A, 6A, and S2), while most limbs were folded (Fig. 4A). Fast and slow walking were classified qualitatively according to the magnitude of trunk undulation (Fig. 4B,C), yielding angular ranges of 31 ± 3 and 12 ± 5 deg, respectively (Figs 5B,C and 6A).

**Table 1.** Summary of locomotor modes and data used for kinematic and EMG analyses.

| Species | Locomotion mode | Number of individuals<sup>*</sup> | Number of trials<sup>*</sup> | Number of recording sites<sup>†</sup> | EMG data note<sup>‡</sup> | Kinematic phase reference |
|---|---|---|---|---|---|---|
| *Scolopendra mutilans* | swimming | 3 | 7 | 13 | #3 right only | trunk |
|  | fast walking | 2 | 6 | 12 |  | trunk, limb |
|  | slow walking | 2 | 4 | 8 |  | limb |
| *Scolopocryptops rubiginosus* | swimming | 3 | 9 | 12 | #3 excluded | trunk, limb |
|  | fast walking | 3 | 9 | 9 | #2 left only, #3 excluded | limb |
|  | slow walking | 1 | 3 | 6 |  | limb |

<sup>\*</sup>Number of individuals and trials included in the kinematic analyses. <sup>†</sup>Number of electromyogram (EMG) recording sites retained after recording-quality screening. <sup>‡</sup>Numbers following # identify individuals within each species.

**Figure 4.**
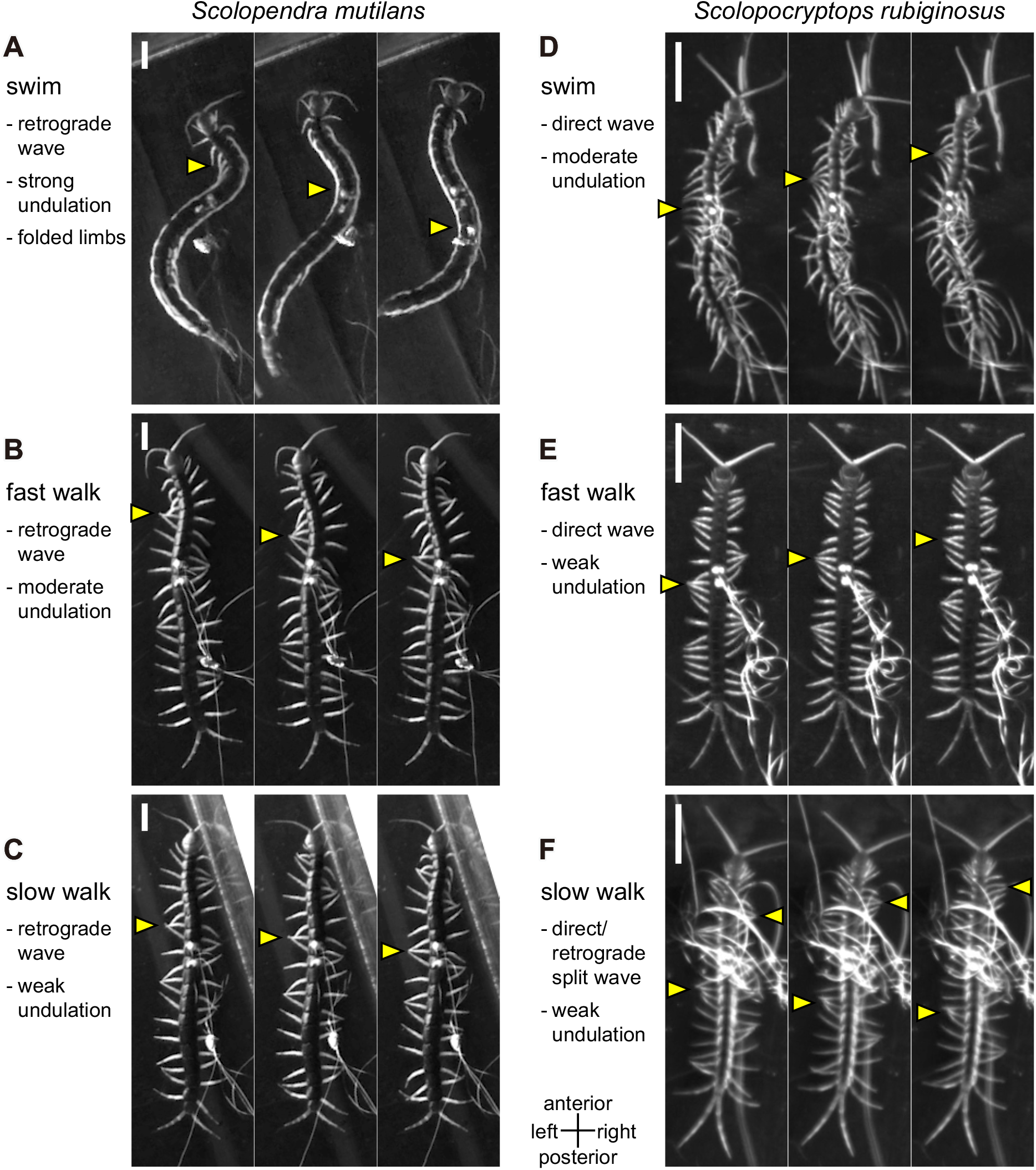
Dorsal-view snapshots during experiments in *Scolopendra mutilans* (A–C) and *Scolopocryptops rubiginosus* (D–F). (A,D) Swimming. (B,E) Fast walking. (C,F) Slow walking. Arrowheads indicate representative regions with limb crowding and/or concave trunk bending. The three panels for each species show the same individual. In each panel, the field of view remains fixed across snapshots. Time progresses from left to right, with intervals of 0.2 s in (A,C) and 0.1 s in the others. Electrode wires are visible, particularly in (F). The corresponding videos are shown in Video S1. Scale bars: 1 cm.

**Figure 5.**
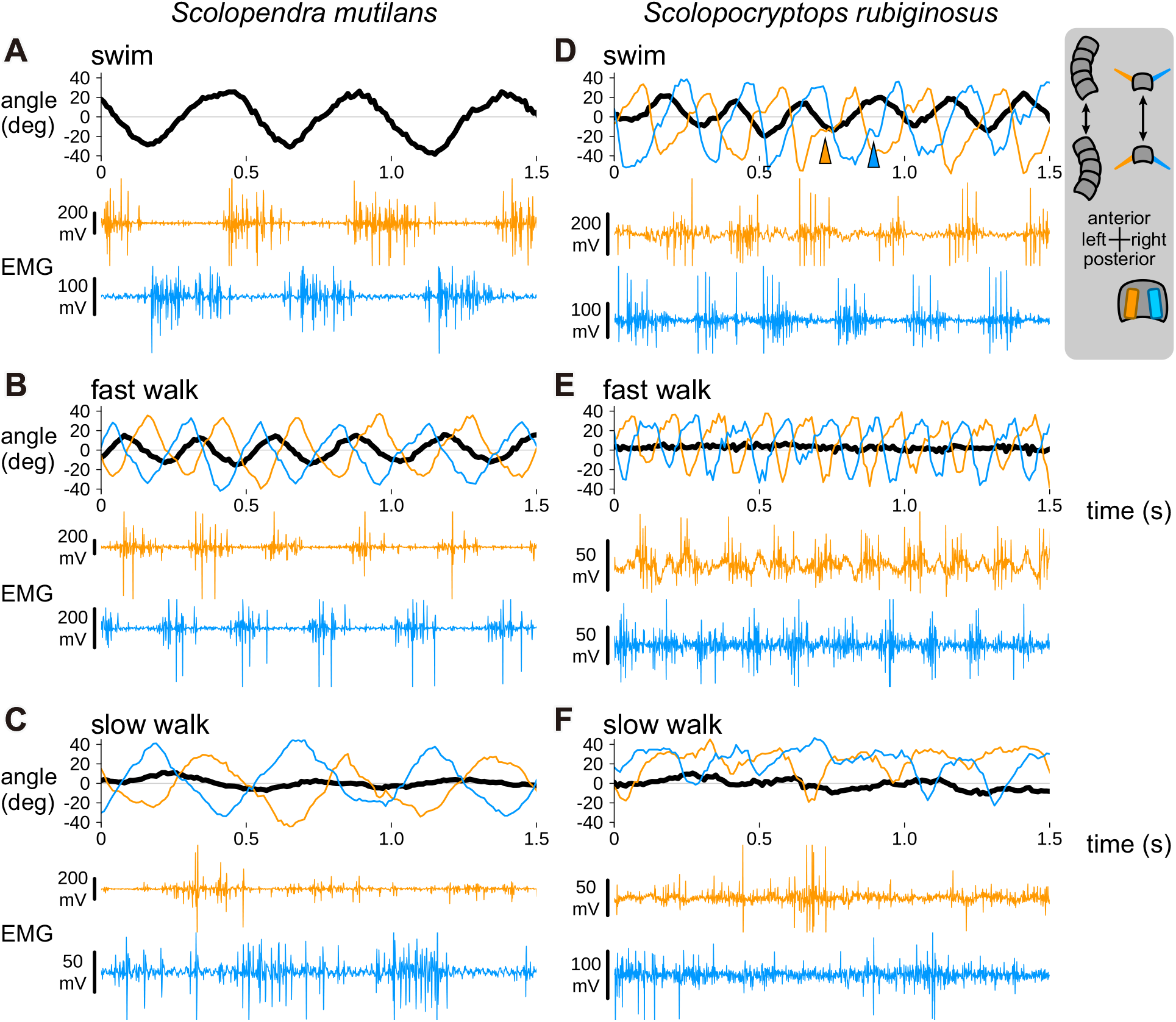
Time series of kinematics and electromyograms (EMGs) in *Scolopendra mutilans* (A–C) and *Scolopocryptops rubiginosus* (D–F). (A,D) Swimming. (B,E) Fast walking. (C,F) Slow walking. The traces were obtained from the same trials as those shown in Fig. 5. Trunk angles were calculated from segments 8, 10, and 12, whereas limb angles and EMGs were obtained from segment 10 (cf. Fig. 2D). In the angle plots, thick black lines represent trunk angles, while orange and light blue lines represent left and right limb angles, respectively. Limb angles are not shown in (A), where the limbs were folded. Arrowheads in (D) indicate representative instances of relatively prolonged movement from the maximally posterior to the mid-forward limb position. In the EMG plots, orange and light blue traces represent signals recorded from the left and right deep dorsal oblique muscles (DDOMs), respectively. Schematics at the upper right illustrate the positive and negative directions of trunk and limb angles and the bilateral EMG recording sites. The three panels for each species show the same individual. Each panel shows a 1.5-s excerpt from the 2-s trial shown in the corresponding panel of Fig. 4.

In *S. rubiginosus*, swimming was characterized by trunk undulation with an angular range of 35 ± 13 deg (Figs 5D and 6A) without limb folding; both the trunk and limbs exhibited direct-wave patterns (Fig. 4D). Walking with a fully direct wave was classified as fast walking (Fig. 4E). Walking with a mixed pattern resembling the split pattern described by Kuroda et al., 2022, with a direct wave in the anterior region and a retrograde wave in the posterior region, was classified as slow walking (Fig. 4F). Trunk undulation was weak during both fast and slow walking, with angular ranges of 8 ± 3 and 15 ± 4 deg, respectively (Figs 5E,F and 6A).

**Figure 6.**
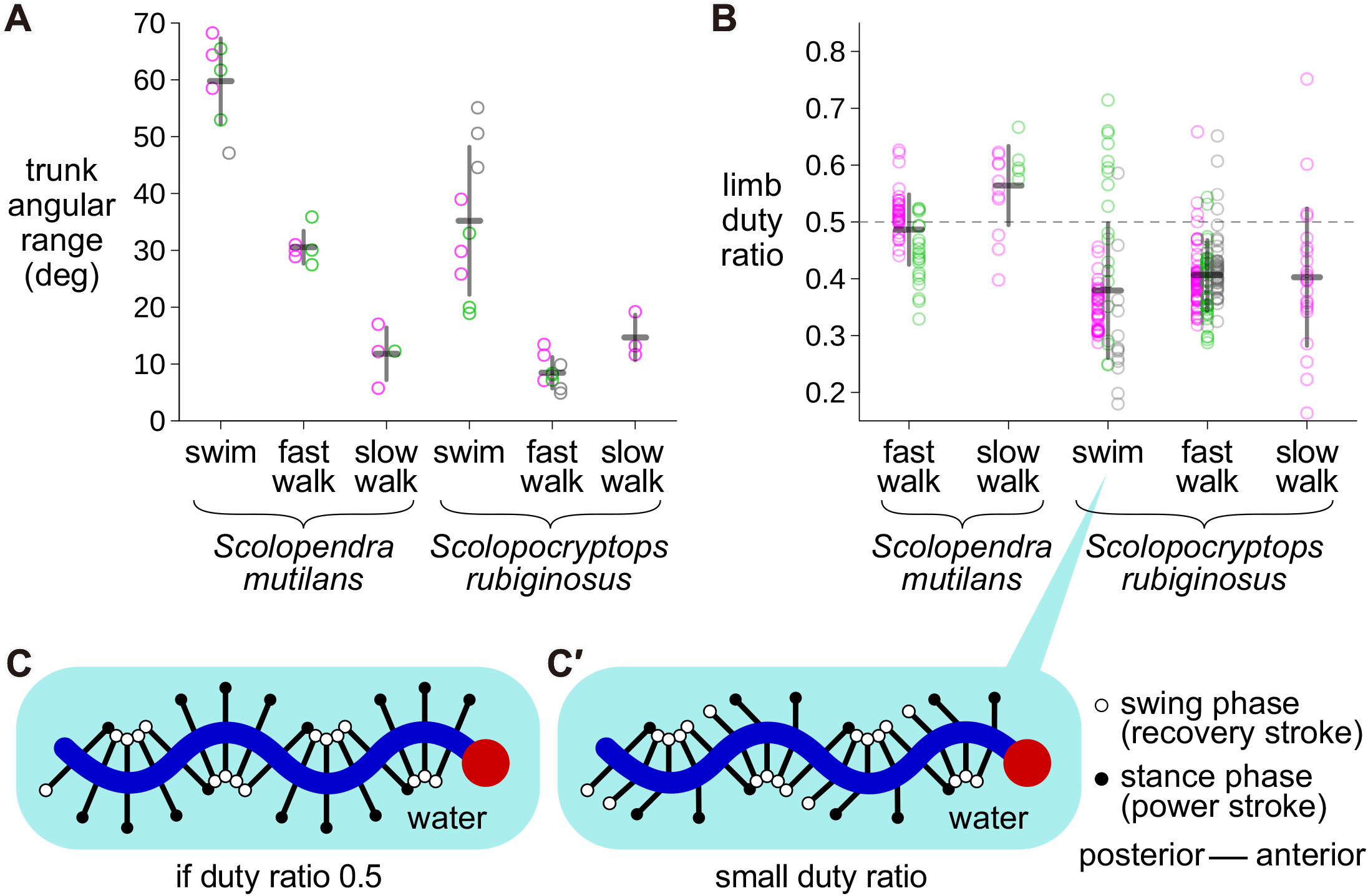
Kinematic characteristics. (A) Angular range of trunk undulation. Each point represents one 2-s trial. (B) Limb duty ratio. Each point represents one limb cycle, with data from the left and right limbs pooled. The dashed line indicates a duty ratio of 0.5. In (A,B), colors distinguish individuals, horizontal bars indicate the mean, and vertical bars indicate SD. (C) Hypothetical swimming scheme assuming a duty ratio of 0.5, using the same schematic representation as in Fig. 1E. (C′) Updated swimming scheme for *Scolopocryptops rubiginosus* based on the present results, with a smaller duty ratio of approximately 0.4, indicative of a relatively rapid power stroke.

The limb duty ratio was 0.49 *±* 0.06 during fast walking and 0.56 *±* 0.07 during slow walking in *S. mutilans* (Fig. 6B). The ratio tended to be smaller in *S. rubiginosus*, being 0.38 *±* 0.12 during swimming, 0.41 ± 0.06 during fast walking, and 0.40 ± 0.12 during slow walking. Inspection of the limb kinematic waveforms indicated that the movement from the maximally posterior to the mid-forward position (phase 0 to *π/*2) was relatively prolonged (Fig. 5D, arrowheads).

### Kinematic relationships among the trunk and limbs

We examined trunk–limb phase relationships during fast walking in *S. mutilans* and swimming in *S. rubiginosus*, in which both trunk and limb kinematics showed clear periodicity. During fast walking in *S. mutilans*, the trunk–limb phase difference remained close to 0 across the four representative limb phases (Fig. 7A). Thus, the maximally anterior and posterior limb positions approximately coincided with straight trunk postures, whereas the mid-forward and mid-backward limb positions corresponded to maximal convex and concave trunk bending, respectively.

**Figure 7.**
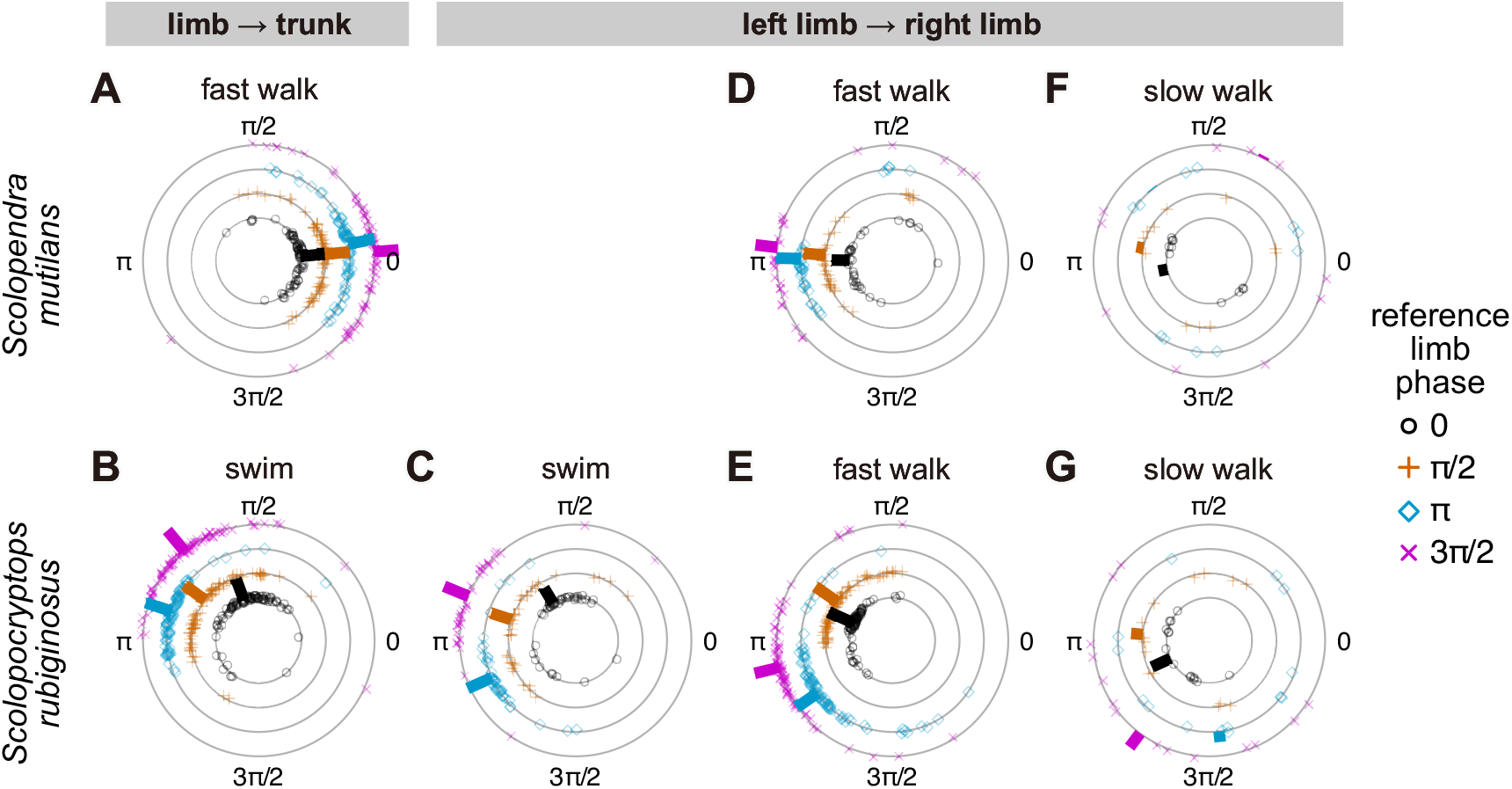
Phase relationships among trunk and limb kinematics. Upper row: *Scolopendra mutilans*; lower row: *Scolopocryptops rubiginosus*. (A,D,E) Fast walking. (B,C) Swimming. (F,G) Slow walking. In (A,B), the limb was used as the reference, and the phase difference between the trunk and limb phases was calculated at the four representative limb phases (0, *π/*2, *π*, and 3*π/*2). Trunk phase was defined separately for the left and right sides so that maximal convex bending toward the limb corresponded to phase *π/*2, and the left- and right-side data were pooled (cf. Fig. 3). In (C–G), the left limb was used as the reference, and the phase difference between the right and left limb phases was calculated at the four representative left-limb phases. Symbols indicate the representative phases used for each comparison: open circles, 0; plus signs, *π/*2; diamonds, *π*; and crosses, 3*π/*2. Angular position represents the phase difference wrapped to the range 0–2*π*. The four concentric circles distinguish the four reference phases. Thick bars indicate the circular mean phase difference, with bar length representing phase consistency (mean resultant length).

During swimming in *S. rubiginosus*, the trunk–limb phase difference was generally around 3*π/*4 but varied among the four representative limb phases (Fig. 7B), indicating that the phase relationship was not constant throughout the locomotor cycle. The kinematic waveforms showed temporal asymmetry between the recovery and power strokes (Fig. 5D, arrowheads). In particular, the limb was close to the lateral position near peak trunk bending during the recovery stroke, whereas it remained more posterior during the power stroke. Likewise, the trunk tended to be nearly straight when the limb reached its anterior extreme but remained bent when the limb reached its posterior extreme. This asymmetry was consistent with the relatively short power stroke indicated by the smaller limb duty ratio.

We next examined contralateral limb coordination using the left limb as the reference. During swimming in *S. rubiginosus*, the phase difference between the right and left limbs varied among the four representative phases of the left limb (Fig. 7C). In particular, the phase difference when the left limb was maximally posterior (phase 0) deviated substantially from *π*. This variation was consistent with the temporal asymmetry of limb movement associated with the relatively rapid power stroke.

During fast walking, *S. mutilans* exhibited an approximately antiphase relationship between the left and right limbs, with little variation among the four representative phases (Fig. 7D). In contrast, the phase difference in *S. rubiginosus* remarkably changed within the locomotor cycle (Fig. 7E). During slow walking, phase differences between the left and right limbs were more variable, and no clear contralateral phase relationship was observed in either *S. mutilans* (Fig. 7F) or *S. rubiginosus* (Fig. 7G).

### Phase-dependent trunk muscle activity

We mapped EMG intensity onto kinematic phases in each condition, using the kinematics of the body part that showed clear periodicity (Table 1, Fig. 5). In trunk-based phase analyses, trunk muscle activity showed similar phase-dependent patterns during swimming in both species and fast walking in *S. mutilans* (Fig. 8A–C). EMG activity was intense from before the convex bending peak (*π*/2) to before the concave bending peak (3*π*/2) of the trunk cycle. The cosinor analysis yielded preferred phases of µ = 0.87*π*, 0.83*π*, and 0.69*π*, with amplitudes of *A* = 1.21, 0.67, and 1.22, for swimming in *S. mutilans*, swimming in *S. rubiginosus*, and fast walking in *S. mutilans*, respectively. In *S. mutilans*, the preferred phase during fast walking was slightly earlier than that during swimming.

**Figure 8.**
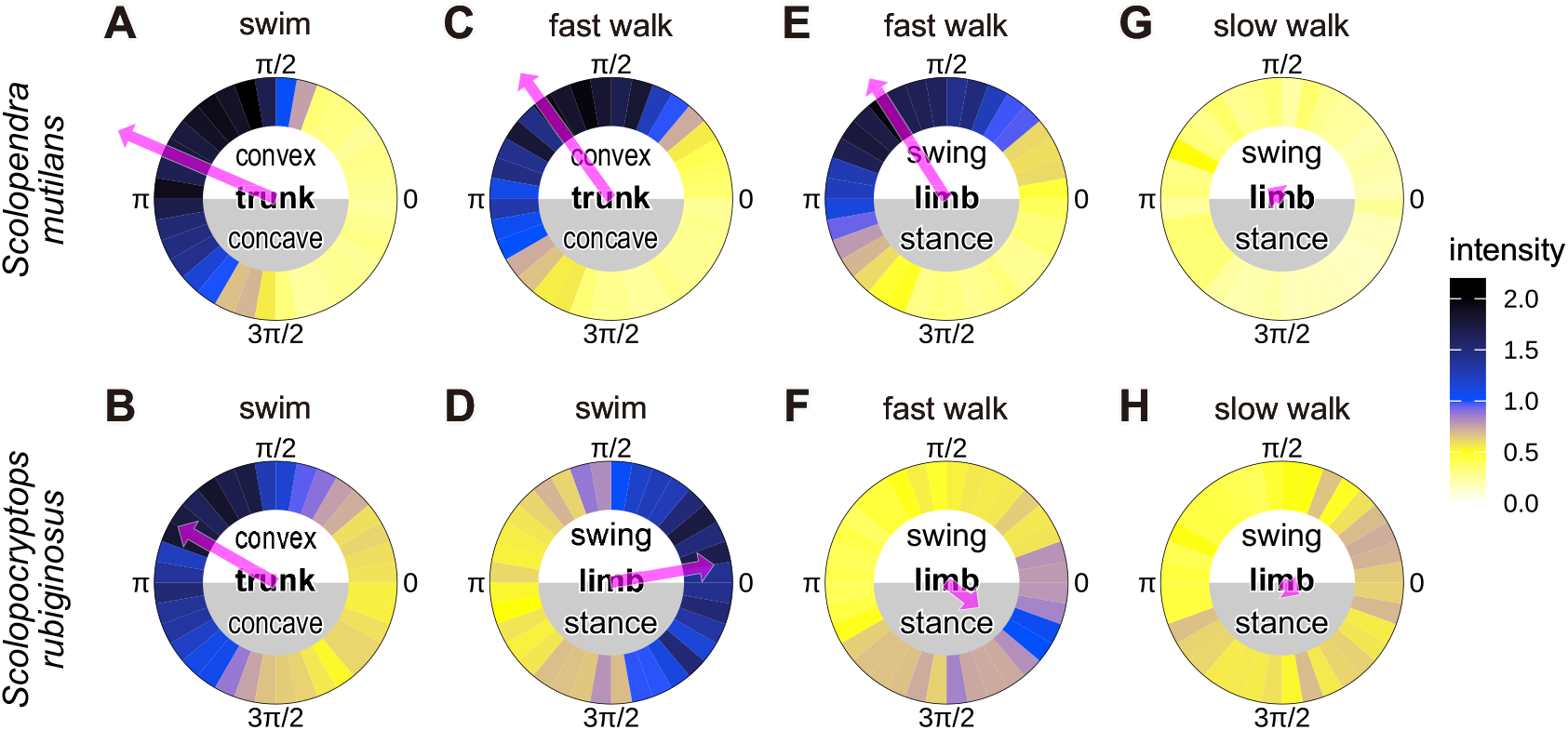
Electromyogram (EMG) intensity mapped onto kinematic phases. Upper row: *Scolopendra mutilans*, lower row: *Scolopocryptops rubiginosus*. In (A–C), EMG intensity was mapped onto the phase of trunk bending defined for the EMG recording side (cf. Fig. 3A). In (D–H), EMG intensity was mapped onto the phase of the limb on the EMG recording side (cf. Fig. 3B). (A,B,D) Swimming, (C,E,F) fast walking, (G,H) slow walking. The arrow in each circle represents the preferred phase with its direction and length corresponding to the preferred direction (µ) and the amplitude (*A*).

In limb-based phase analyses, EMG activity during swimming in *S. rubiginosus* was elevated around the maximally posterior limb position (phase 0), with a preferred phase of *µ* = 0.04*π* and an amplitude of *A* = 0.62 (Fig. 8D). During fast walking, the timing of EMG activity differed between the retrograde-wave gait of *S. mutilans* and the direct-wave gait of *S. rubiginosus* (Fig. 8E,F). In *S. mutilans*, EMG activity was intense from after the beginning of the swing phase (0) to after that of the stance phase (*π*) (*µ* = 0.66*π, A* = 1.11; Fig. 8E). In contrast, *S. rubiginosus* showed weaker phase-dependent variation, with relatively higher EMG intensity from after the beginning of the stance phase (*π*) to after that of the swing phase (0) (*µ* = 1.78*π, A* = 0.24; Fig. 8F). During slow walking, EMG intensity showed little phase-dependent variation in both species (Fig. 8G,H). The preferred phases were *µ* = 0.82*π* (*A* = 0.16) in *S. mutilans* and *µ* = 1.80*π* (*A* = 0.12) in *S. rubiginosus*.

## DISCUSSION

### Contralateral limb coordination in relation to trunk undulation

We discuss kinematics in this section and its relationships with EMG in later sections. We examined kinematic phase relationships across locomotor modes with different degrees of trunk undulation.

In scolopendromorph centipedes, left and right limbs have been reported to move in antiphase during straight locomotion (Manton, 1952). Such contralateral coordination can contribute to net forward propulsion despite strong lateral trunk undulation. Consistent with this interpretation, the mean phase differences between the left and right limbs were close to *π* in *Scolopocryptops rubiginosus* swimming and *Scolopendra mutilans* fast walking (Fig. 7C,D). For the left and right limbs to remain in antiphase throughout the cycle, the swing and stance phases must be equally long (limb duty ratio: 0.5). This condition was approximately met during fast walking in *S. mutilans* (Fig. 6B), consistent with the nearly constant antiphase relationship observed throughout the cycle (Fig. 7D).

However, during swimming with direct-wave trunk undulation in *S. rubiginosus*, the mean limb duty ratio was approximately 0.4 despite strong trunk undulation (Fig. 6B). The absence of substrate contact during swimming may have allowed a rapid power stroke and a slower recovery stroke, with the limbs reaching their posterior positions early and remaining near their lateral positions for longer during phases 0–*π*/2 (Fig. 5D, arrowheads). This asymmetry may explain two features of the observed phase relationships. First, contralateral limb phase differences vary within the cycle (Fig. 7C), resulting in asymmetric limb angles at the peak of trunk bending, with the limb on the concave side oriented laterally and that on the convex side directed posteriorly (Fig. 6C′), rather than symmetric limb angles (Fig. 6C). Second, the rapid power stroke may advance the limb phase relative to the trunk phase, reducing their phase difference to approximately 3*π/*4 (Fig. 7B, circles and crosses), compared with the *π* expected from the opposite wave direction to *S. mutilans* (Fig. 7A). During the slower recovery stroke, the phase difference may return toward *π* as the limb approaches its anterior position (Fig. 7B, diamonds). The prolonged recovery stroke to match limb movements with the trunk cycle may involve phase resetting, in which the phase of a locomotor rhythm is reset in response to sensory feedback or perturbations, as suggested by physiological and mathematical studies in insects and mammals (Büschges, 1995; Schomburg et al., 1998; Rybak et al., 2006; Ambe et al., 2021).

In contrast, when trunk undulation is weak, straight locomotion does not necessarily require a fixed phase relationship between the left and right limbs. In our kinematic analysis, the mean left–right limb phase difference was close to *π* during fast walking in *S. rubiginosus* (Fig. 7E), whereas no strong phase relationship was observed during slow walking in *S. mutilans* (Fig. 7F). This contrast may be related to trunk muscle activity, as discussed later.

### Common trunk muscle activation: *S. mutilans* swimming vs. *S. rubiginosus* swimming

We now turn to an integrated discussion of the kinematic and EMG findings. We begin with trunk-based phase analyses, comparing limb-folded retrograde swimming in *S. mutilans* and limbed direct swimming in *S. rubiginosus* (Fig. 8A,B). In both species, EMG activity was consistently elevated when the recorded side of the trunk was convex and reduced as it became concave. Since each trunk segment bilaterally accommodates a pair of DDOMs (Fig. 2D), contraction on one side of the trunk pulls that side toward a concave curvature, as suggested by Manton (1965) and experimentally supported by Anderson et al. (1995). This mechanism explains the swimming kinematics of *S. mutilans* (Fig. 8A), in which trunk undulation is primarily generated by trunk muscle activity, as the limbs are folded against the trunk. This mechanical role of trunk muscle contraction, which converts convex curvature into concave curvature on the active side, is consistent with that of axial muscles in vertebrates such as snakes and lampreys (Jayne, 1988; Williams et al., 1989).

In contrast, swimming in *S. rubiginosus* involves direct trunk undulation accompanied by limb strokes, yet the phase relationship between trunk muscle activity and trunk bending was nearly identical to that observed during swimming in *S. mutilans*, which exhibits retrograde undulation without limb strokes (Fig. 8A,B). This indicates that the relationship between trunk muscle activity and trunk kinematics is maintained regardless of whether the limbs contribute to propulsion or the direction in which the trunk undulation propagates. Furthermore, this similarity suggests that trunk undulation during swimming in *S. rubiginosus* is actively generated by trunk muscles rather than arising merely as a passive reaction to limb movements.

### Effects of limb movements on trunk muscle activation: *S. mutilans* swimming vs. fast walking

We next compare trunk-based phase relationships during retrograde swimming without limb strokes and retrograde fast walking with limb strokes in *S. mutilans* (Fig. 8A,C). Trunk muscle activity occurred earlier during fast walking, with EMG activity beginning before the recorded side of the trunk became convex. We consider three possible explanations for this difference. First, limb movements may induce passive changes in trunk kinematics during fast walking. Second, the difference can arise from the locomotor cycle period. Although phase-normalized data do not retain absolute temporal information, the frequency of fast walking was approximately twice that of swimming (Fig. 5A,B). If the absolute time delay between the onset of muscle activity and the resulting kinematic deformation remains constant, the delay expressed as a fraction of the locomotor cycle would be larger during fast walking. Third, neural coupling between trunk and limb movements may be involved. Delvolvé et al. (1997) reported that trunk muscle activity in limb-bearing regions of newts preceded that in more anterior regions and proposed neural coupling between limb and trunk locomotor systems. This interpretation is consistent with our observation that trunk muscle activity occurred earlier in the locomotor cycle when limb movements were involved.

The temporal delay between trunk muscle activity and kinematics during fast walking warrants further consideration. In the trunk-based phase analysis, the timing of EMG onset indicates a delay between the initiation of muscle activity that produces concave bending and the actual onset of bending toward the concave side (after phase *π*/2; Fig. 8A–C). Similarly, the timing of EMG offset indicates a delay between the cessation of muscle activity and the onset of bending toward the convex side (after phase 3*π*/2). Such delays can arise from the viscoelastic and inertial properties of the trunk, as generally considered in studies of axial locomotion (Frolich and Biewener, 1992), and a similar tendency was observed in our centipedes. Anderson et al. (1995) reported that EMG onset occurred approximately 5% of a locomotor cycle after maximal convex trunk bending in *Scolopendra heros* and regarded these events as nearly coincident. In contrast, our data indicate that EMG onset preceded maximal convex trunk bending by approximately *π*/4 (12.5% of a cycle). One possible explanation for this difference is the location of the recording segment. Anderson et al. (1995) averaged measurements across multiple trunk segments, but their plots suggest variation in the phase relationship among segments, including some in which kinematic changes lagged behind EMG activity. Our results may therefore reflect a delay specific to the 10th trunk segment. A study of salamander locomotion also reported that the delay between trunk muscle activity and bending increased toward the posterior body (Frolich and Biewener, 1992), supporting the possibility that such delays vary among trunk segments.

### Kinematic interpretation of EMG timing: *S. rubiginosus* swimming vs. *S. mutilans* fast walking

In the next three sections, we examine trunk muscle activity relative to limb-based kinematic phases. Because we did not directly measure neural feedback activity, interpretations concerning the presence and direction of neural feedback between the limbs and trunk remain speculative in the three sections.

We first compare direct swimming in *S. rubiginosus* (Fig. 8D) and retrograde fast walking in *S. mutilans* (Fig. 8E), both involving limb movements and trunk undulation. The EMG pattern in Fig. 8D was shifted approximately *π*/4 later than expected from a *π* rotation of that in Fig. 8E, likely reflecting the trunk–limb phase relationships discussed in the preceding section on kinematic coordination. In *S. rubiginosus*, the trunk–limb phase relationship deviated by approximately *π*/4 from that expected based on a *π* shift of the relationship observed in *S. mutilans* (Fig. 7A,B). This deviation can be explained by the limb duty ratio being less than 0.5 during swimming in *S. rubiginosus* (Fig. 6B).

### Contrasting mechanical roles of trunk muscles: *S. mutilans* fast walking vs. *S. rubiginosus* fast walking

Using limb-based phase analyses, we next compare fast walking with strong retrograde trunk undulation in *S. mutilans* and direct fast walking with weak trunk undulation in *S. rubiginosus* (Fig. 8E,F). DDOM activity was strong during the ipsilateral swing phase in *S. mutilans* (Fig. 8E), whereas weak activity was observed during the ipsilateral stance phase in *S. rubiginosus* (Fig. 8F). This difference in the timing of muscle activity relative to the swing and stance phases reflects the opposite directions of the locomotor waves.

Regarding trunk bending, DDOMs in *S. rubiginosus* exhibited weak activity during the stance phase despite the nearly straight trunk during fast walking (Fig. 8F). Since the DDOMs of the 10th trunk segment are connected to the 11th segment, we consider the local mechanical relationship between these two segments. For example, when the left limb of segment 10 pushes against the substrate during the stance phase, the resulting reaction force would tend to passively bend the region between segments 10 and 11 into a left-convex curvature (Fig. 9A). However, contraction of the left DDOM at this timing would generate a force opposing this passive bending, potentially maintaining a straight configuration between the two segments (Fig. 9B). If such active straightening occurs across all trunk segments, it could explain the nearly straight trunk observed during fast walking in *S. rubiginosus*.

**Figure 9.**
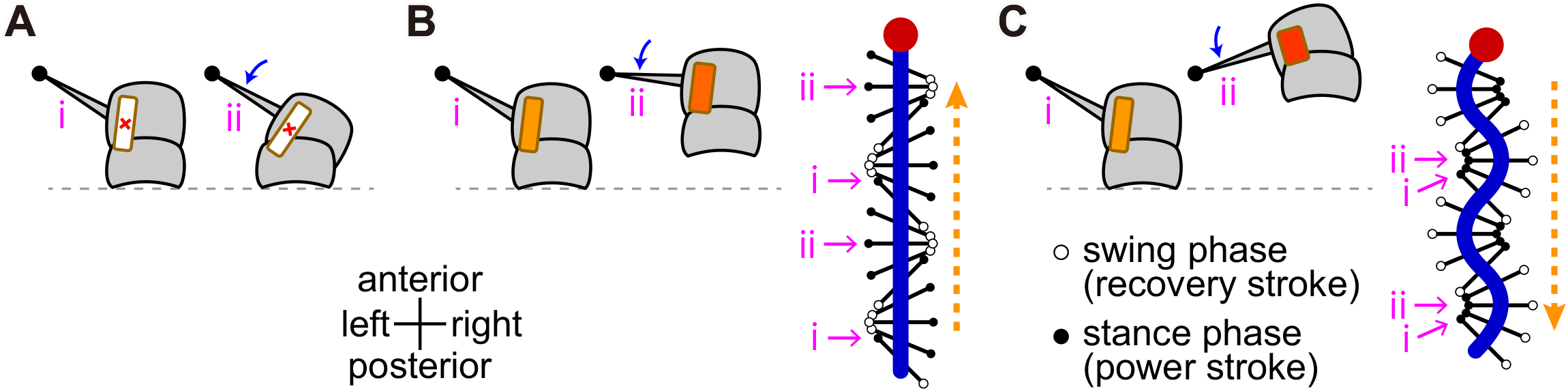
Schematic illustration of the proposed effects of deep dorsal oblique muscle (DDOM) contraction during fast walking. The segment-level illustrations show trunk segments 10 and 11, with only the left limb and DDOM depicted. In each panel, the supporting limb pushes backward from state i to state ii; the curved arrows show the pushing direction. (A) Without DDOM contraction. (B) With weak DDOM contraction, corresponding to straight direct-wave locomotion in *Scolopocryptops rubiginosus*. (C) With strong DDOM contraction, corresponding to undulatory retrograde-wave locomotion in *Scolopendra mutilans*. In (B,C), representative limb positions corresponding to states i and ii are also shown at multiple locations along the body. The dashed arrows indicate the directions of metachronal waves.

Considering only local mechanics, stronger or more prolonged DDOM contraction could further bend the trunk into a left-concave curvature, as observed during fast walking in *S. mutilans* (Fig. 9C). This interpretation, in which DDOM activity actively promotes concave bending rather than merely resisting it, is consistent with that of Anderson et al. (1995). Slow walking in *S. mutilans* represents a mode of walking with reduced trunk bending, as slight trunk bending was occasionally observed (Fig. 5C). Because these interpretations are limited to local trunk–limb mechanics, the different wave directions in Fig. 9B and C may involve mechanisms beyond differences in DDOM contraction strength.

A potentially analogous role of axial muscles in active stabilization has been reported in vertebrates. EMG studies in horses have suggested that the longissimus dorsi muscles primarily function to stabilize the vertebral column against dynamic forces during locomotion (Licka et al., 2004, 2009). Our results suggest that centipede trunk muscles can similarly counteract forces generated during limb propulsion, suggesting a similar stabilizing role of trunk muscles across phylogenetically distant taxa.

### Effects of trunk muscle acticity on limb coordination: *S. rubiginosus* fast walking vs. *S. mutilans* slow walking

Finally, we compared contralateral limb coordination and the dependence of trunk muscle activity on limb-based kinematic phases during fast walking in *S. rubiginosus* and slow walking in *S. mutilans*, both of which involved weak trunk bending. In *S. rubiginosus* fast walking, the left and right limbs moved approximately in antiphase (Fig. 7E), accompanied by phase-dependent trunk muscle activity (Fig. 8F). In contrast, neither a clear contralateral phase relationship (Fig. 7F) nor strong phase-dependent trunk muscle activity (Fig. 8G) was observed in *S. mutilans* slow walking. These findings raise the possibility that the neural circuits underlying periodic trunk muscle activity also contribute to contralateral limb coordination, even when trunk bending is weak. Based on kinematic observations of millipede walking, numerical simulations using a neuromechanical model have shown that local trunk–limb interactions mediated by sensory feedback can generate interlimb coordination without direct coupling among limbs (Yasui et al., 2026). In cats, electrophysiological recordings from trunk and limb motor nerves during fictive locomotion revealed coordinated rhythmic activity, indicating that their coordination can arise from central neural mechanisms (Koehler et al., 1984). In salamanders, EMG recordings combined with trunk and limb kinematic analyses have suggested that periodic trunk muscle activity during walking resists limb-induced forces and may modulate body stiffness, potentially improving motor coordination (Bennett et al., 2001). Further studies examining the causal relationships between trunk muscle activity and limb coordination would help clarify the role of trunk–limb coupling in locomotion.

### Summary: mode-dependent trunk–limb interactions

Our findings suggest that the roles of trunk muscles (DDOMs) and their relationships with limb movements vary across locomotor conditions:

- **During swimming**, without ground reaction forces, trunk muscles actively generate trunk bending regardless of limb involvement in propulsion or wave propagation direction. This is supported by the nearly identical EMG–trunk bending phase relationships in *S. mutilans* without limb strokes and *S. rubiginosus* with limb strokes (Fig. 8A,B). In *S. rubiginosus* swimming, the relatively rapid power stroke (Fig. 6B,C′) can explain why contralateral limb phase differences change within the limb cycle (Fig. 7C).
- **During walking with strong trunk bending**, i.e., *S. mutilans* fast walking, trunk muscle activity can actively promote concave trunk bending (Fig. 9C). The activity begins earlier relative to trunk bending than during swimming (Fig. 8A,C), which may reflect differences in mechanical effects of limb propulsion, locomotor frequency, or neural coupling between trunk and limb movements. The nearly constant antiphase relationship between contralateral limbs (Fig. 7D) can facilitate straight locomotion despite strong trunk bending.
- **During walking with weak trunk bending**, phase-dependent trunk muscle activity may be associated with contralateral limb coordination:
- **With phase-dependent trunk muscle activity**, as in *S. rubiginosus* fast walking (Fig. 8F), trunk muscle activity can resist passive bending induced by limb forces, actively maintaining a nearly straight trunk (Fig. 9B). This alternating activity may contribute to contralateral limb coordination, with the limbs moving approximately in antiphase (Fig. 7E).
- **Without clear phase-dependent trunk muscle activity**, as in *S. mutilans* slow walking (Fig. 8G), contralateral limb coordination may be less constrained, with the limbs showing highly variable phase differences (Fig. 7F).

### Limitations and future directions

Several limitations should be considered when interpreting our results. First, EMG recordings were restricted to DDOMs at a single trunk segment. Simultaneous recordings from multiple successive segments could reveal how muscle activation propagates along the body and help explain the mechanisms underlying the differences between direct and retrograde undulation. Second, walking was examined under tethered conditions on a slippery substrate, where limb movements occurred without forward displacement (Video S1). The mechanical interactions between the limbs, trunk, and substrate can therefore differ from those during natural terrestrial locomotion, in which the limbs obtain strong traction against the ground.

Although our results revealed differences in the timing of trunk muscle activity relative to limb movements between direct and retrograde locomotion, the mechanisms producing these differences remain unresolved. Additional EMG recordings from limb muscles could characterize their activation patterns but would not, by themselves, establish the neural interactions between limb and trunk control. Direct recordings from interneuronal circuits involved in trunk–limb coordination would help clarify the presence and direction of neural coupling. Alternatively, a biomechanical model incorporating active muscle contraction, passive trunk mechanics, and the timing of muscle activation and mechanical responses could provide a constructive explanation for the observed coordination patterns.

Overall, our findings suggest that trunk muscles actively generate bending or resist limb-induced deformation depending on the locomotor mode, while their periodic neural activation may also be associated with contralateral limb coordination. In particular, the nearly straight trunk during fast walking in *S. rubiginosus* can be explained by active stabilization against limb-induced forces (Fig. 9B). Understanding locomotor control in multisegmented animals therefore requires considering not only how muscles generate movement, but also how they stabilize the body and potentially contribute to limb co-ordination. Our study highlights that the mechanisms underlying locomotor coordination cannot always be inferred from observable body movements alone.

## ACKNOWLEDGMENTS

Animal collection was conducted with the support of Seto Marine Biological Laboratory, Kyoto University. We thank Dr. Emily Standen (University of Ottawa) and Dr. Shura Suzuki (Tohoku University) for helpful discussions, and Dr. Shunsuke Shigaki (National Institute of Informatics) for assistance with the experimental setup.

## ETHICS

Ethical approval was not required for this study because centipedes are not subject to animal ethics approval requirements under the applicable institutional and national guidelines.

## DATA ACCESSIBILITY

The kinematic data, EMG data, and analysis code are provided in the electronic supplementary material, Data S1. Representative locomotor recordings are provided in Video S1.

## DECLARATION OF AI USE

We used ChatGPT (OpenAI) to assist with English language editing and the development of code for data analysis. The authors reviewed and validated the AI-assisted code and take full responsibility for the analyses and manuscript content.

## AUTHORS’ CONTRIBUTIONS

D.W.: conceptualization, data curation, formal analysis, funding acquisition, investigation, methodology, project administration, software, supervision, validation, visualization, writing–original draft, writing– review and editing; K.Y.: conceptualization, data curation, funding acquisition, investigation, methodology, project administration, resources, supervision, validation, visualization, writing–original draft, writing–review and editing. Both authors gave final approval for publication and agreed to be held accountable for the work performed therein.

## SUPPLEMENTARY MATERIAL

**Figure S1.**
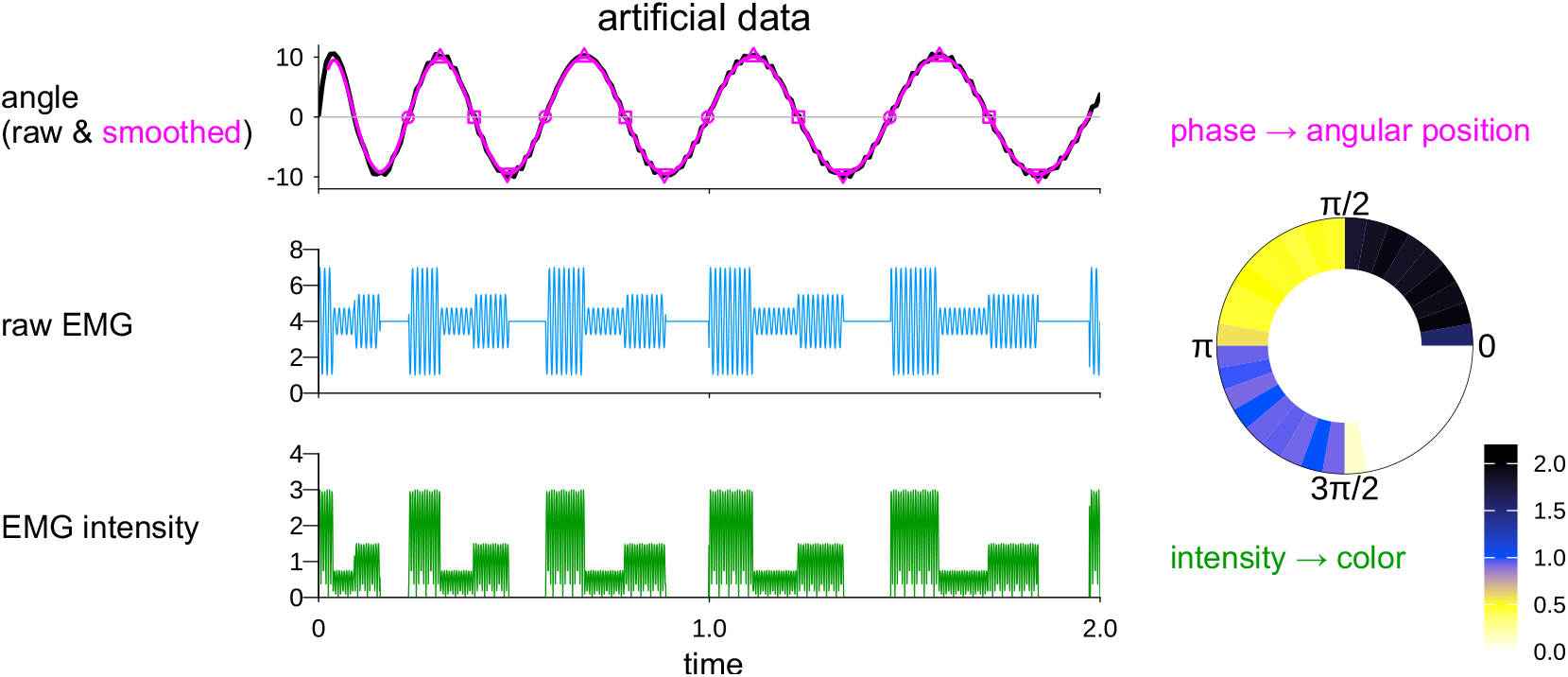
Schematic illustration of kinematic phase extraction and Electromyogram (EMG) intensity mapping using artificial data. The raw angle trace (black) was smoothed (magenta). In this figure, the trunk phase was defined assuming a left-side EMG recording (cf. Fig. 3A, left): circles, upward zero crossings (phase 0); upward triangles, positive peaks (*π/*2); squares, downward zero crossings (*π*); and downward triangles, negative peaks (3*π/*2). The raw EMG signal (blue) was median-centered, and its absolute value was taken to obtain EMG intensity (green). The resulting EMG intensity was averaged within each kinematic phase bin and visualized in a circular plot (right), where angular position represents kinematic phase and color represents EMG intensity.

**Figure S2.**
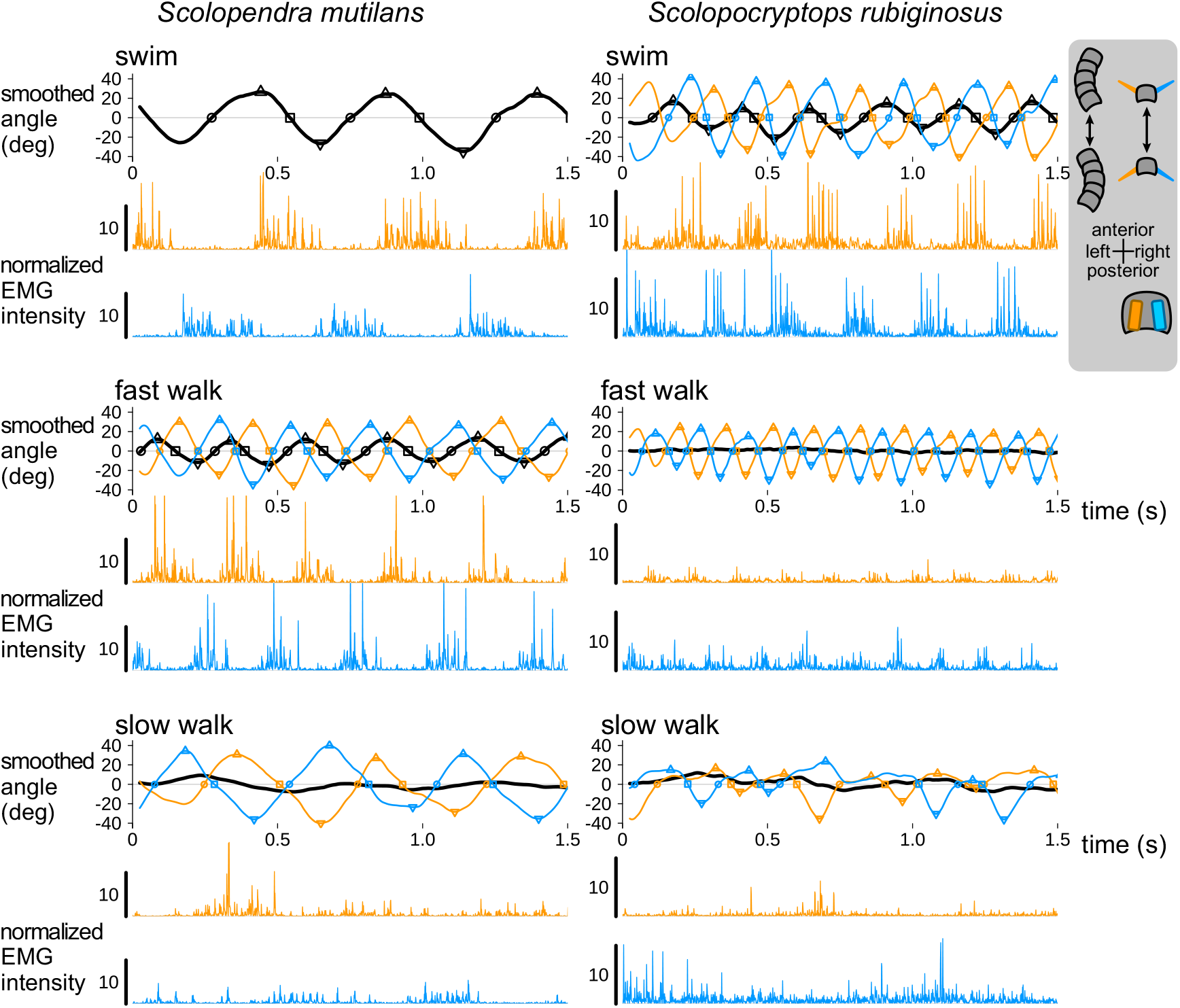
Time series of processed kinematics and electromyogram (EMG) intensity in *Scolopendra mutilans* (left column) and *Scolopocryptops rubiginosus* (right column). Swimming, fast walking, and slow walking are shown from top to bottom. The traces were obtained from the same trials as those shown in Fig. 5. In the angle plots, thick black lines represent trunk angles, while orange and light blue lines represent left and right limb angles, respectively. Trunk and limb angle traces were mean-centered within each trial and smoothed using a running mean filter. Symbols indicate representative points used for phase extraction: circles, upward zero crossings; upward triangles, positive peaks; squares, downward zero crossings; and downward triangles, negative peaks. In the EMG plots, orange and light blue traces represent normalized intensity recorded from the left and right deep dorsal oblique muscles (DDOMs), respectively. EMG signals were median-centered within each trial, rectified, and normalized by the largest mode-specific mean intensity for each individual and recording side.

**Video S1. Representative locomotor modes in two centipede species**. Dorsal-view recordings of swimming, fast walking, and slow walking in *Scolopendra mutilans* and *Scolopocryptops rubiginosus* are shown sequentially. Each video clip shows a 2-s (400-frame) trial corresponding to a panel in Fig. 4. Scale bars: 1 cm.

**Data S1. Kinematic and electromyographic (EMG) data and analysis code**. The ZIP archive contains kinematic and EMG data for each trial in CSV format, together with the R Markdown code used for the analyses (code.Rmd, readable with any text editor).

## REFERENCES

Ambe, Y., Aoi, S., Tsuchiya, K., and Matsuno, F. (2021). Generation of direct-, retrograde-, and sourcewave gaits in multi-legged locomotion in a decentralized manner via embodied sensorimotor interaction. Frontiers in Neural Circuits, 15:706064.

Anderson, B. D., Shultz, J. W., and Jayne, B. C. (1995). Axial kinematics and muscle activity during terrestrial locomotion of the centipede scolopendra heros. Journal of Experimental Biology, 198(5):1185–1195.

Bennett, W. O., Simons, R. S., and Brainerd, E. L. (2001). Twisting and bending: the functional role of salamander lateral hypaxial musculature during locomotion. Journal of Experimental Biology, 204(11):1979–1989.

Brown, D., Christian, W., and Hanson M R. (2025). Tracker video analysis and modeling tool. https://physlets.org/tracker/, Accessed: Dec. 2, 2025.

Büschges, A. (1995). Role of local nonspiking interneurons in the generation of rhythmic motor activity in the stick insect. Journal of neurobiology, 27(4):488–512.

Delvolvé, I., Bem, T., and Cabelguen, J.-M. (1997). Epaxial and limb muscle activity during swimming and terrestrial stepping in the adult newt, pleurodeles waltl. Journal of Neurophysiology, 78(2):638–650.

Frolich, L. M. and Biewener, A. A. (1992). Kinematic and electromyographic analysis of the functional role of the body axis during terrestrial and aquatic locomotion in the salamander ambystoma tigrinum. Journal of Experimental Biology, 162(1):107–130.

Jayne, B. C. (1988). Muscular mechanisms of snake locomotion: An electromyographic study of lateral undulation of the florida banded water snake (nerodia fasciata) and the yellow rat snake (elaphe obsoleta). Journal of Morphology, 197:159–181.

Koehler, W. J., Schomburg, E. D., and Steffens, H. (1984). Phasic modulation of trunk muscle efferents during fictive spinal locomotion in cats. The Journal of Physiology, 353(1):187–197.

Kuroda, S., Uchida, N., and Nakagaki, T. (2022). Gait switching with phase reversal of locomotory waves in the centipede scolopocryptops rubiginosus. Bioinspiration & Biomimetics, 17(2):026005.

Lewis, J. G. E. (1981). The Biology of Centipedes. Cambridge University Press.

Licka, T., Frey, A., and Peham, C. (2009). Electromyographic activity of the longissimus dorsi muscles in horses when walking on a treadmill. The Veterinary Journal, 180(1):71–76.

Licka, T. F., Peham, C., and Frey, A. (2004). Electromyographic activity of the longissimus dorsi muscles in horses during trotting on a treadmill. American journal of veterinary research, 65(2):155–158.

Manton, S. M. (1952). The evolution of arthropodan locomotory mechanisms. spart 3. the locomotion of the chilopoda and pauropoda. Zoological Journal of the Linnean Society, 42(284):118–167.

Manton, S. M. (1965). The evolution of arthropodan locomotory mechanisms. part 8. functional requirements and body design in chilopoda, together with a comparative account of their skeleto-muscular systems and an appendix on a comparison between burrowing forces of annelids and chilopods and its bearing upon the evolution of the arthropodan haemocoel. Zoological Journal of the Linnean Society, 45(306–307):251–484.

Rybak, I. A., Shevtsova, N. A., Lafreniere-Roula, M., and McCrea, D. A. (2006). Modelling spinal circuitry involved in locomotor pattern generation: insights from deletions during fictive locomotion. The Journal of physiology, 577(2):617–639.

Schomburg, E., Petersen, N., Barajon, I., and Hultborn, H. (1998). Flexor reflex afferents reset the step cycle during fictive locomotion in the cat. Experimental brain research, 122(3):339–350.

Williams, T. L., Grillner, S., Smoljaninov, V. V., Wallén, P., Kashin, S., and Rossignol, S. (1989). Locomotion in lamprey and trout: The relative timing of activation and movement. Journal of Experimental Biology, 143:559–566.

Yasui, K., Kano, T., Standen, E. M., Aonuma, H., Ijspeert, A. J., and Ishiguro, A. (2019). Decoding the essential interplay between central and peripheral control in adaptive locomotion of amphibious centipedes. Scientific reports, 9(1):18288.

Yasui, K., Yamaichi, T., and Ishiguro, A. (2026). Millipedes suggest decentralized trunk–limb coordination control is essential for inter-limb coordination in many-legged locomotion. Artificial Life and Robotics, 31(1):34–45.

